# Programmable DNA Nanocages Enhance Levodopa Delivery for Neuroprotection in a Zebrafish Model of Parkinson’s Disease

**DOI:** 10.64898/2026.08.04.742731

**Authors:** Krupa Kansara, Shubha Lakshmi Swain, Bhagyesh Parmar, Shaivee Chokshi, Keya Jantrania, Ashutosh Kumar, Dhiraj Bhatia

## Abstract

Parkinson’s Disease (PD) is the second-most prevalent neurodegenerative disease, often characterized by neural motor dysfunction, oxidative stress, and dopamine receptor malfunction leading to improper dopamine levels in the system. DNA tetrahedron nanostructures are a promising drug delivery agent due to their biocompatibility and properties of controlled and sustained release. In this study we evaluated the potential of using TD-mediated Levodopa delivery for a MPTP induced Parkinson’s Disease model in Zebrafish larvae. The induction of Parkinsonism led to morphological behaviour changes like the presence of tremors, erratic swimming behaviour, latency, and reduced locomotor activity, even elevated reactive oxygen species (ROS) and apoptosis was observed. These effects and symptoms were alleviated when the larvae were treated using TD:Levodopa conjugates, particularly at the 1:100 ratio. At the molecular level, genes like TH, DAT, SOX2, PARKIN and apoptotic genes like BCL2 and caspases showed alteration in expression in the Parkinsonism model and post treatment was induced. This highlights the potential of using DNA nanocages as a novel drug delivery agent as therapeutic strategy for Parkinson’s disease.

**TOC:** Dopamine loaded DNA nanocages with the capacity to overcome biological barriers for release of dopamine with neuroprotection activity in Parkinson’s disease model of zebrafish.

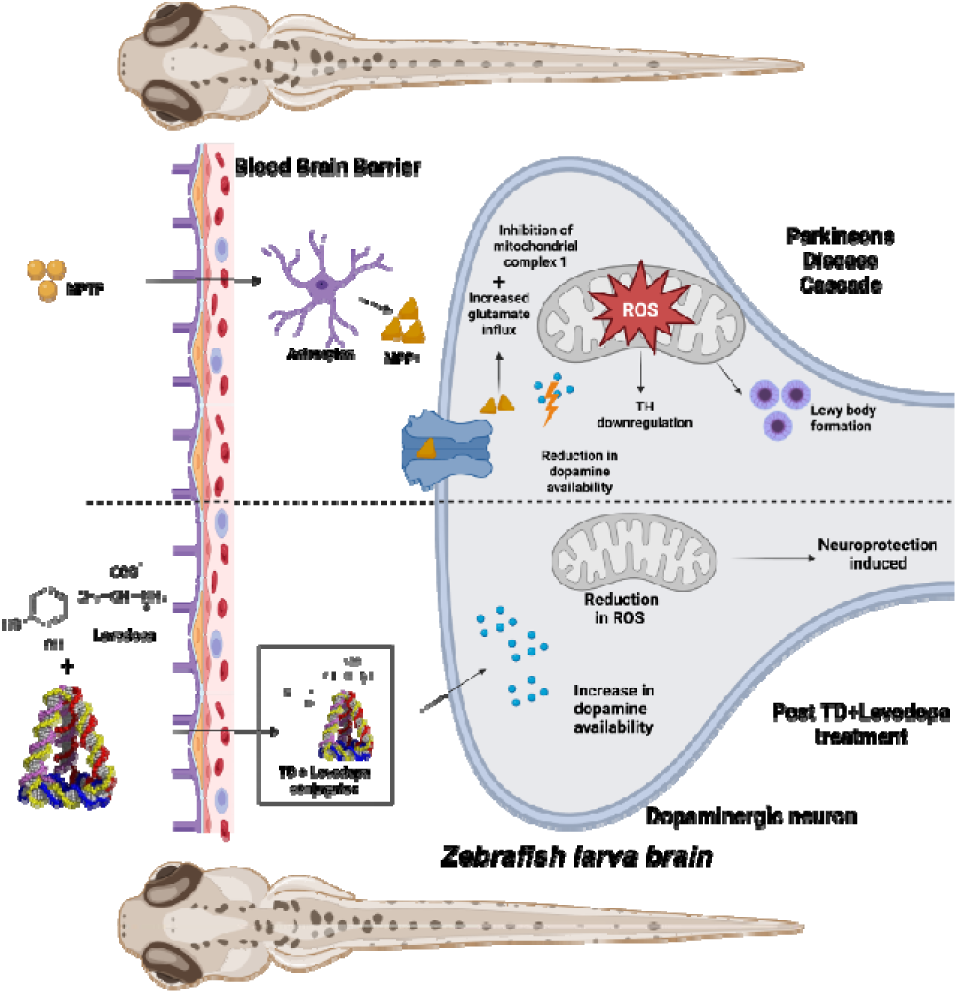

## 1 Introduction

Parkinson’s Disease (PD) is the second-most prevalent neurodegenerative disease[1]. PD occurs due to the progressive degeneration of dopaminergic neurons in the substantia nigra, further leading to the three cardinal symptoms of PD-static tremor, muscle rigidity, and bradykinesia[1, 2]. PD can manifest both motor and non-motor symptoms; the latter often precedes the motor symptoms by several years. The non motor symptoms include hyposomnia because of uncoordinated mouth and throat movement, sleep disorders, depression, autonomic dysfunction, mood disorders, cognitive impairment, gastrointestinal symptoms, and pain, thereby affecting the overall quality of life[1, 3, 4]. The origin of PD involves a complex interaction between genetic predisposition and environmental influences, with age being the most prominent risk factor. Epidemiological research has consistently demonstrated substantial differences in disease prevalence across geographical regions and populations, underscoring the roles of both genetic and environmental factors[3]. Treatment of PD is largely focused on symptom management rather than curing PD. Pharmacological, non-pharmacological, and surgical treatments remain the common conventional treatment approaches for PD. Apart from these, there are new emerging and experimental approaches for treatment[5]. When medical treatments become insufficient, management of PD can be done through non-drug therapies, exercise therapies, psychological interventions, and surgical options[6]. Levodopa, a precursor of dopamine, is the gold standard pharmacological approach widely used for treating PD. To increase its effectiveness and reduce any potential side effects levodopa is often combined with carbidopa. Other pharmacological treatment options include dopamine agonists, monoamine oxidase B (MAO-B) inhibitors, and catechol-O-methyltransferase (COMT) inhibitors. The main goal of these pharmacological treatments is to mimic or enhance dopamine activity in the brain. Non-pharmacological treatment options are used as an adjuvant therapy; on the other hand, the surgical therapy is only advised for the PD patients of advanced stage[5, 6]. However, some of the patients often experience fluctuating symptoms, side effects due to medicines, and disease progression[7, 8]. As a result, a novel therapeutic approach that can achieve controlled release, facilitate easy penetration across the blood brain barrier (BBB), and exhibit lower off-target effects is needed to overcome the issues faced by conventional therapy.

Among pharmacological treatment options, Levodopa remains the gold standard for PD. Several notable studies have demonstrated the effectiveness of Levodopa in the treatment of Parkinson’s disease, including the ELLDOPA and PD MED trials. Findings from these trials indicate that levodopa therapy provides significant symptomatic improvement compared with other dopaminergic therapies, particularly in the early and moderate stages of the disease[9, 10]. Levodopa is a dopamine precursor that can cross the BBB and be converted to dopamine[5]. The therapeutic dose of Levodopa varies among PD patients, although most individuals initially respond well to doses of approximately 300–600 mg/day[11]. Levodopa when administered as monotherapy, only 1-3% of the drug reaches the brain because of premature metabolization and incomplete BBB passage. To increase levodopa’s bioavailability to the brain and reduce oral dosage, levodopa is combined with a peripheral decarboxylase inhibitor (PDI), such as carbidopa or benserazide. These inhibitors inhibit the l-amino acid decarboxylase (AADC) enzyme, thereby preventing premature levodopa metabolization[12].

The underlying mechanism of PD comprises several linked cellular pathways, ultimately leading to the death of dopaminergic neurons in the substantia nigra[3]. Intracellular homeostasis of α-synuclein is maintained via the ubiquitin–proteasome and lysosomal autophagy systems. Accumulation of α-synuclein is a characteristic hallmark of PD and results from impaired pathways due to oxidative stress, mitochondrial dysfunction, or neuronal inflammation. Mitochondrial dysfunction, exacerbated by mutations in genes such as LRRK2, DJ-1, Parkin, and PINK1, can lead to increased neuronal death. The exacerbation of neuronal damage in PD occurs because oxidative stress and neuroinflammation are closely interlinked, thereby reinforcing each other[13]. As a result, neuroprotective therapies that enhance neuronal resistance to oxidative stress and mitochondrial dysfunction, when used parallelly with pharmacological treatments such as Levodopa, have the potential to slow dopaminergic neuronal loss and potentially modify disease progression in Parkinson’s disease. In case of neuronal damage, nanomaterials have been used as neuroprotective agents. The use of nanotechnology as a potential therapeutic for brain injury is still in its early stages, owing to the complexity of the nervous system and potential neurotoxicity of nanomaterials. Previous studies on DNA tetrahedrons indicate that they exert significant biological effects, including neuroprotection and anti-inflammatory activity, owing to their efficient synthesis and good biocompatibility, determined by their DNA composition[14]. TDs have been widely used in the field of bioscience[15, 16], notably in neuromedicine[17, 18] due to their advantages. According to several studies, TDs can promote the proliferation, migration, and neuronal differentiation of neural stem cells (NSCs) and also precisely modulate autophagy[19]. A study by Zhou et.al, found that synergistic therapy with TDs and NSCs improved recovery from spinal cord injury.[20] Owing to their benefit of crossing the blood brain barrier, these nanocages are very useful in treating neurological disorders. It is essential to overcome the blood-brain barrier for drugs to reach brain tissue. This can be achieved by using TDs, since they have been proven useful for delivering various bioactive molecules[21]. Yang and colleagues demonstrated that TDs could cross the BBB to some extent in normal nude mice[22]. Several studies have demonstrated TdN uptake in zebrafish embryos[17, 18, 23–25].

In this study, TDs were designed and synthesized by self-hybridization of four single-stranded oligonucleotides to form a compact three-dimensional tetrahedral structure. To the best of our knowledge, levodopa (3,4-dihydroxy-L-phenylalanine), a clinically proven gold standard treatment for Parkinson’s disease, was conjugated to the TDs framework to create TD: Levodopa nanoconjugates for the very first time, leading to the increased therapeutic effect against MPTP induced parkinsonism as a novel conjugate. The goals of this study were to enhance the stability of Levodopa in a physiological setting, improve its pharmacokinetic profile, reduce the drug dosage, and facilitate targeted delivery across biological barriers. To assess the neuroprotective role of these nanostructures, a zebrafish (*Danio rerio*) model of PD was developed. Owing to their notable genetic and physiological similarity to humans, optical transparency during early development, and suitability for real-time imaging and behaviour analysis, zebrafish have been recognized as a robust vertebrate model for the study of neurodegenerative diseases. TDs and TD:Levodopa nanoconjugates were used to treat Parkinson-like symptoms induced by a chemical neurotoxin. Post-treatment evaluations comprised morphological and behavioral analysis, and quantification assays. Additionally, it documents detailed biological response profiles in zebrafish models of Parkinson’s disease, providing an essential baseline for future translational studies of DNA nanostructure-based drug delivery systems for neurodegenerative disorders.

## 2 Materials and methods

### 2.1 Revealing the Distinctive Features of TDs and Formulated TD:Levodopa

#### 2.1.1 Synthesis of TDs and TD:Levodopa Conjugates

A previously described one-pot synthesis method was used to synthesize TD nanostructures[26]. Four complementary single-stranded DNA oligos (M1, M2, M3, and M4) were combined in an equimolar concentration with 2 mM MgCl_2_, followed by a thermal annealing procedure performed in a PCR apparatus with cycling settings from 95 to 4 °C. After 10 minutes of heating to 95 °C, the reaction mixture was progressively cooled to 4 °C. Oligonucleotides labelled with cyanine-5 (Cy5) were employed for imaging. To functionalise the TD:Levodopa conjugates, a combination of TD and Levodopa in a 1:100 molar ratio was incubated for 5 hours.

#### 2.1.2 Characterisation and conjugation of TDs and TDs with Levodopa

##### DLS and Zeta Potential

DLS and zeta potential analyses were performed using a Malvern Analytical Zetasizer Nano-ZS to determine the hydrodynamic size and charge distribution of TDs and TD:Levodopa conjugations (1:100). Following a Gaussian fit, the data were plotted using GraphPad Prism software.

##### Atomic Force Microscopy (AFM)

Atomic Force Microscope (AFM) was used to analyse morphological characteristics and structural integrity of the self-assembled TDs and TD: Levodopa conjugations (1:100). Sample preparation was done as per the established lab protocols. Aliquots (5-10 µl) of TD and TD:Levodopa (1:100) were spread onto a freshly sliced mica substrate to obtain the AFM pictures in tapping mode with RFESPG-75 (antimony-doped Si) and SCANASYST-AIR (silicon nitride) probes in ambient circumstances. The imaging was performed with a Bruker NanoWizard Sense + Bio AFM installed at IIT Gandhinagar, Gujarat.

##### Electrophoretic Mobility Shift Assay

Native-PAGE was used in an electrophoretic mobility shift assay. A 10% polyacrylamide gel was used to assess the formation of the TD and TD:Levodopa (1:100). After 90 minutes of operation at 90 V, the gel was stained with EtBr and examined using the Bio-Rad ChemiDoc MP imaging system for gel documentation.

### 2.2 Determination of Levodopa entrapment efficiency in TDs and kinetic release

The TD:Levodopa (1:100) were stored inside the dialysis membrane, which was then immersed in a beaker containing phosphate buffer solution (PBS) with a pH of 5.5, and 7.4 while being continuously stirred. Post 12-hour incubation, the PBS solution was collected to assess the entrapment efficiency. Subsequently, the PBS was replaced with fresh PBS (pH 5.5, and 7.4), and the setup was maintained for an additional 72 hours to collect hourly samples (1hr to 8hr, 12hr, 24hr, 48hr, and 72 hour) for drug release analysis. The drug content in the collected sample was quantified using a spectrophotometer. Entrapment efficiency was calculated based on the amount of drug released within the first 12 hours, and the drug release kinetics were analysed subsequently.

#### Zebrafish maintenance

An optimum laboratory environment was maintained with 14-hour light/10-hour dark cycles, and 27°C - 28°C lab temperature was maintained. Adult zebrafish were housed in 20-litre aquarium tanks with mildly saline water (Red Sea Coral Pro salt), and constant aeration was provided using pumps. A constant pH of water ranging from 6.8 to 7.2 was maintained and checked using a pH meter (Horiba). The fish were fed brine shrimp (Artemia) twice daily and basic flakes (Aquafin) once daily. A lab-built breeding box was used for breeding, and fish were kept in the ratio of 3:4 (male: female). Maximising egg production was achieved by separating the fish two days before breeding twice weekly. Spawning happened one hour post-removal of dividers. The eggs were kept in a sterile Petri plate comprising E3 medium (5.0 mM NaCl, 0.17 mM KCl, 0.33 mM CaCl₂, and 0.33 mM MgSO₄) and were stored in an incubator at 28 °C until further experimentation. The experiments were conducted on 3 days post-fertilisation (dpf) larvae[27].

#### Induction of Parkinsonism-like symptoms in zebrafish larvae

Healthy larvae of 3 dpf were selected for further experiments. MPTP treatment was given to healthy larvae at a 300 µM concentration for 3 days. The concentration of MPTP was decided based on a literature review and experiments conducted at 250 µM, 300 µM, 350 µM and 500 µM[28]. These larvae were closely monitored for behavioural and morphological changes. Several factors, such as increased latency, increased erratic swimming and tremors, were monitored to visually know if the Parkinson like symptoms had been induced. The time frame for inducing Parkinsonism was fixed at 3 days to ensure minimal mortality. Once induced, the larvae were introduced to three recovery methods; 300 µM levodopa[29] solution made in 5% dextrose[30], levodopa conjugated with TDs and E3 media. The larvae were monitored for 3 hours post-treatment. Selected healthy larvae from the batch were used as a control group.

#### Quantification of Parkinsonism-like symptoms in zebrafish larvae

##### Morphological assessment of PD-induced zebrafish larvae

Ten to fifteen larvae at 3dpf were subjected to MPTP treatment and later to recovery treatments (E3 media, 300 µM levodopa, TDs conjugated with levodopa). Activity, latency, erratic swimming, mortality, and malformations were observed and documented at three time stamps (1 hour, 2 hours, and 3 hours) post-treatment. Heart rate and tremors were measured 3 days post-MPTP treatment and 3 hours after the recovery treatments. These changes were examined using a stereo microscope.

##### Intracellular reactive oxygen species evaluation in PD-induced zebrafish larvae

The oxidative stress in larvae was measured by quantifying the generation of reactive oxidative species (ROS), using DCFDA dye. Concentration of the DCFDA dye was 2mM in a 6-well plate and was incubated for 1 hour at 28℃ in the dark. After incubation, three E3 media washes were given. Imaging was carried out in a fluorescent microscope equipped with a filter at 450nm - 490nm. The fluorescence intensities were measured in ImageJ software[27].

##### Programmed Cell Death evaluation in PD-induced zebrafish larvae

Visualisation of apoptotic cells in the larvae was done using acridine orange dye. Larvae subjected to treatments were taken in a 6-well plate, and the protocol for acridine orange staining was followed. The larvae were exposed to 10 µg/ml of acridine orange stain in E3 media, and the plate was incubated for 30 minutes at 28℃ in the dark. After incubation, the larvae were washed thrice with E3 media. Imaging was carried out in a fluorescent microscope equipped with a filter at 450nm - 490nm. The fluorescence intensities were measured using the ImageJ software[27].

##### Enzyme linked immunosorbent assay (ELISA)

Dopamine protein levels were quantified using the Zebrafish Dopamine ELISA Kit (ELK9307) as per the instruction manual provided by the manufacturer. Standards and samples were added to antibody-coated wells and incubated to allow antigen binding. After the addition of the detection reagent and streptavidin-HRP, TMB substrate was used for colour development. The reaction was stopped using a stop buffer, and absorbance was measured at 450 nm using a microplate reader. Dopamine concentrations were calculated from a standard curve generated with known standards.

##### Detection of Park7/DJ-1 expression in Zebrafish Larvae by Immunofluorescent Assay

In order to find Park7/DJ-1 aggregates using an immunofluorescent test, ten to fifteen larvae at 3dpf were treated with MPTP and then recovered with TD, levodopa, and TD:LD (1:100). The larvae were fixed overnight at 4°C in 4% paraformaldehyde (PFA) made in phosphate-buffered saline (PBS). After fixation, the larvae underwent three PBS washes before being permeabilized for 30 minutes at room temperature using 0.5% Triton X-100 in PBS. Before blocking, a proteinase K treatment (10 µg/mL in PBS) was administered. To reduce non-specific antibody binding, samples were blocked for two hours at room temperature using 5% bovine serum albumin (BSA) made in PBS containing 0.1% Triton X-100. Rabbit anti-PARK7/DJ-1 primary antibody (1:200 dilution in blocking solution) was used to incubate the larvae for an entire night at 4°C. After primary antibody incubation, samples were washed three times with PBS for 10 min each and incubated with Alexa Fluor 594 goat anti-rabbit IgG (1:500; Thermo Fisher Scientific) for 2 h at room temperature in the dark. The larvae were completely cleaned with PBS after the secondary antibody incubation, and their nuclei were seen for 20 minutes by counterstaining them with Hoechst 33342 (1 μg/mL). After final washes, the larvae were mounted in an antifade mounting media on glass slides. Fluorescence images were acquired using a confocal laser scanning microscope with identical laser power, detector gain, and acquisition settings for all experimental groups. Nuclei were visible in the blue channel (Hoechst 33342), while PARK7/DJ-1 immunoreactivity was seen in the red fluorescence channel (Alexa Fluor 594). ImageJ was used to quantitatively analyze PARK7/DJ-1 aggregates, including aggregate-positive area and fluorescence intensity, following background subtraction. To ensure an objective comparison between experimental groups, all pictures were subjected to the identical thresholding parameters, and fluorescence values were normalized to those of the control group.

##### Gene expression studies using qRT-PCR

Total RNA isolation was performed at two-time stamps for the experiment. Initial RNA isolation was performed 3 days post MPTP treatment, followed by RNA isolation for the recovery groups 3 hours post treatment. Favoregen RNA isolation kit was used for RNA isolation. cDNA synthesis was carried out using 1000ng of isolated RNA, which was followed by a real-time PCR. The genes associated with PD and its recovery were assessed, such as TH (tyrosine hydroxylase), DAT (dopamine transporter), Sox2, and Parkin. The apoptotic genes were also assessed, such as Caspase3 (Cysteine-aspartic acid protease 3), Caspase9 (Cysteine-aspartic acid protease 9), and Bcl2 (B-cell Lymphoma 2). GAPDH was used as a housekeeping gene for both cases.

## Results and Discussion

### Characterization and conjugation of TDs and TDs with Levodopa DLS and Zeta Potential

Dynamic Light Scattering (DLS) was used to assess the hydrodynamic size distribution and dispersion of the TDs and TD:Levodopa conjugates. The samples were diluted in a nuclease-free buffer to reduce multiple scattering effects, and the measurements were carried out at 25 °C with a standard scattering angle of 90°. The DLS results were in accordance with the predicted outcomes, as they showed a narrow and monodisperse size distribution with an average hydrodynamic diameter of roughly 11.23±1.56 nm for TDs, and 21±1.26 for TD: Levodopa (1:100). The polydispersity index (PDI) of the samples is also low which suggests that there is a high degree of homogeneity and effective self-assembly without notable aggregation or multimodal particle populations. Zeta potential was used to measure the surface charge and colloidal stability of the TdN and TdN: Levodopa (1:100). It was observed that the zeta potential for the TDs was −10.32 ±1.22 mV, which is in agreement with the structure, due to the negatively charged phosphate backbone of DNA. The zeta potential for TD:Levodopa (1:100) was recorded at −3.31 ±1.22 mV, which showed successful conjugation of Levodopa molecules on the TDs. The combined analysis of DLS and zeta potential measurements confirms the successful synthesis of uniform, stable, and evenly distributed TDs, as well as their conjugation with Levodopa. The negative zeta potential in aqueous media is crucial for ensuring colloidal stability and preventing particle aggregation through electrostatic repulsion. Furthermore, these findings, in conjunction with AFM observations, provide robust confirmation of the TDs’ effective synthesis and structural integrity.

### Atomic Force Microscopy (AFM)

Atomic Force Microscopy (AFM) topography showed uniformly dispersed nanostructures, specifically TDs and TD:Levodopa (1:100) complexes, exhibiting distinct triangle or dot-like characteristics. The measured lateral dimensions, ranging from 12 to 15 nm, closely matched the predicted size based on the specified DNA model. The observed slight variations in apparent size are attributed to partial flattening on the mica substrate and tip convolution effects. Height profile analysis confirmed the three-dimensional tetrahedral configuration, revealing an average height of 5-7 nm. The homogeneity and monodispersity of the nanostructures suggest high assembly fidelity and structural integrity. Furthermore, the absence of noticeable aggregation or deformation demonstrated the stability and durability of the self-assembled TDs and TD:Levodopa (1:100) under imaging conditions. The AFM analysis successfully confirmed the creation of distinct, uniform TDs and TD:Levodopa(1:100) nanocages with structural traits consistent with the intended architecture. These results validate the accuracy of the DNA-based construction at the nanoscale and support the effectiveness of the assembly procedure.

### Electrophoretic Mobility Shift Assay

The TDs and TD:Levodopa (1:100) were assembled sequentially, as shown by native polyacrylamide gel electrophoresis. In order to confirm the successful hybridization and structure formation, the electrophoretic mobility of individual oligonucleotides, partial assemblies, and complete TDs was analyzed. Distinct mobility alterations were observed as the assembly of DNA strands began. It was observed that single-stranded oligonucleotides moved faster, and bands were seen at the bottom of the gel. Decreased electrophoretic mobility was observed in the intermediate bands representing partially formed structures, indicating a higher molecular weight. The fully completed TDs showed a single characteristic band with the slowest migration, showcasing the formation of a compact, higher-order tetrahedral structure. Faint bands of TD:Levodopa (1:100) were observed due to the conjugation of the drug over TDs.

**Figure 1:**
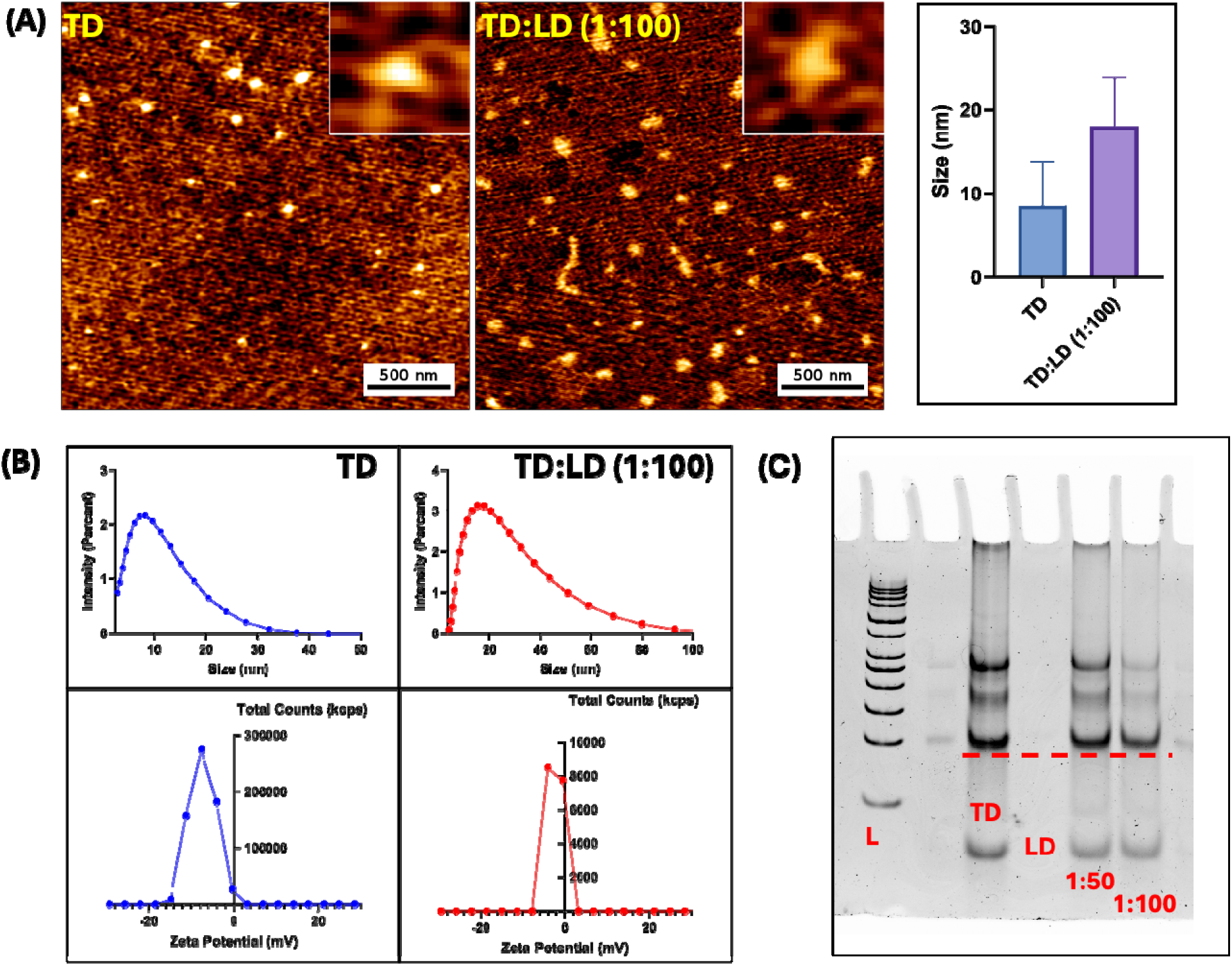
Characterization of a tetrahedral DNA nanostructure A) AFM Images of TDs and TD:Levodopa conjugates, scale bar is 500nm B) DLS results of the synthesized tetrahedron showed the hydrodynamic diameter and Zeta potential of negatively charged DNA nanoparticles C) Gel electrophoresis mobility shift-based characterization showing the formation of the Tetrahedron and TDN: Levodopa (1:50), (1:100).

### Levodopa entrapment efficiency in TDs and kinetic release

Entrapment efficiency is known as the percentage of the formulation’s total levodopa that is successfully encapsulated within the TDs. The entrapment efficiency of the drug within the TDs was found to be 99.70% and 99.78% in pH 7.4 and 5.5, respectively. The drug was not tested at the basic pH due to its properties, as it degraded at a basic pH. This indicates that the drug effectively binds with the drug carrier without undergoing degradation and assists in sustained release at the target site. It was observed in the release kinetics that at pH 7.4, a sustained release is observed, which increases over time, indicating that the treatment is effective for a longer time period, rather than a rapid and abrupt release. Similarly, in an acidic pH, the release is sustained and increases over time, assisting in targeted delivery at the compromised site. Targeted delivery ensures that the drug is not released elsewhere, affecting other tissues and cells, protecting them from the blanket effect of the drug.

**Figure 2:**
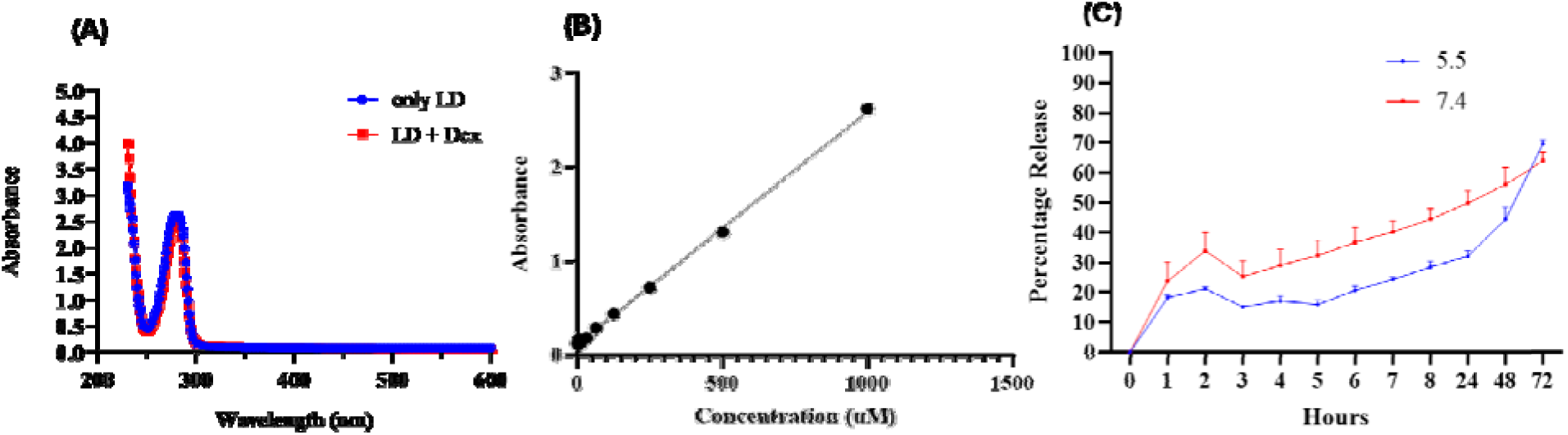
In vitro release kinetics of Levodopa from DNA Tetrahedrons at pH 5.5 and 7.4. (A) Absorbance spectra of Levodopa with and without dextrose (B) Standard curve of Levodopa (C). The graph shows the cumulative percentage of Levodopa released over time at pH 5.5 and 7.4. Data represent means ± SEM of three independent experiments.

### Morphological and behavioural observations

The physiological effects of Levodopa, TDs, and TD:Levodopa conjugates were evaluated through detailed morphological and behavioural assessment of zebrafish larvae following treatment. Key parameters, including locomotor activity, latency, erratic swimming, malformations, and mortality, were monitored under both normal and induced Parkinsonism. During the initial hours following treatment, measurable physiological alterations were recorded over a 3-hour observation period. At 1 hour post-treatment, locomotor activity was minimal across experimental groups, reflecting a temporary suppression of normal neuromotor function in the early stages following Parkinson’s induction. Detectable activity at this stage was observed only in the TD and TD:Levodopa (1:100) groups, suggesting an early onset of functional recovery in treatments involving DNA tetrahedron as a delivery agent. As the observation period progressed, a gradual restoration of locomotor behaviour became evident across the treatment groups.

**Figure 3:**
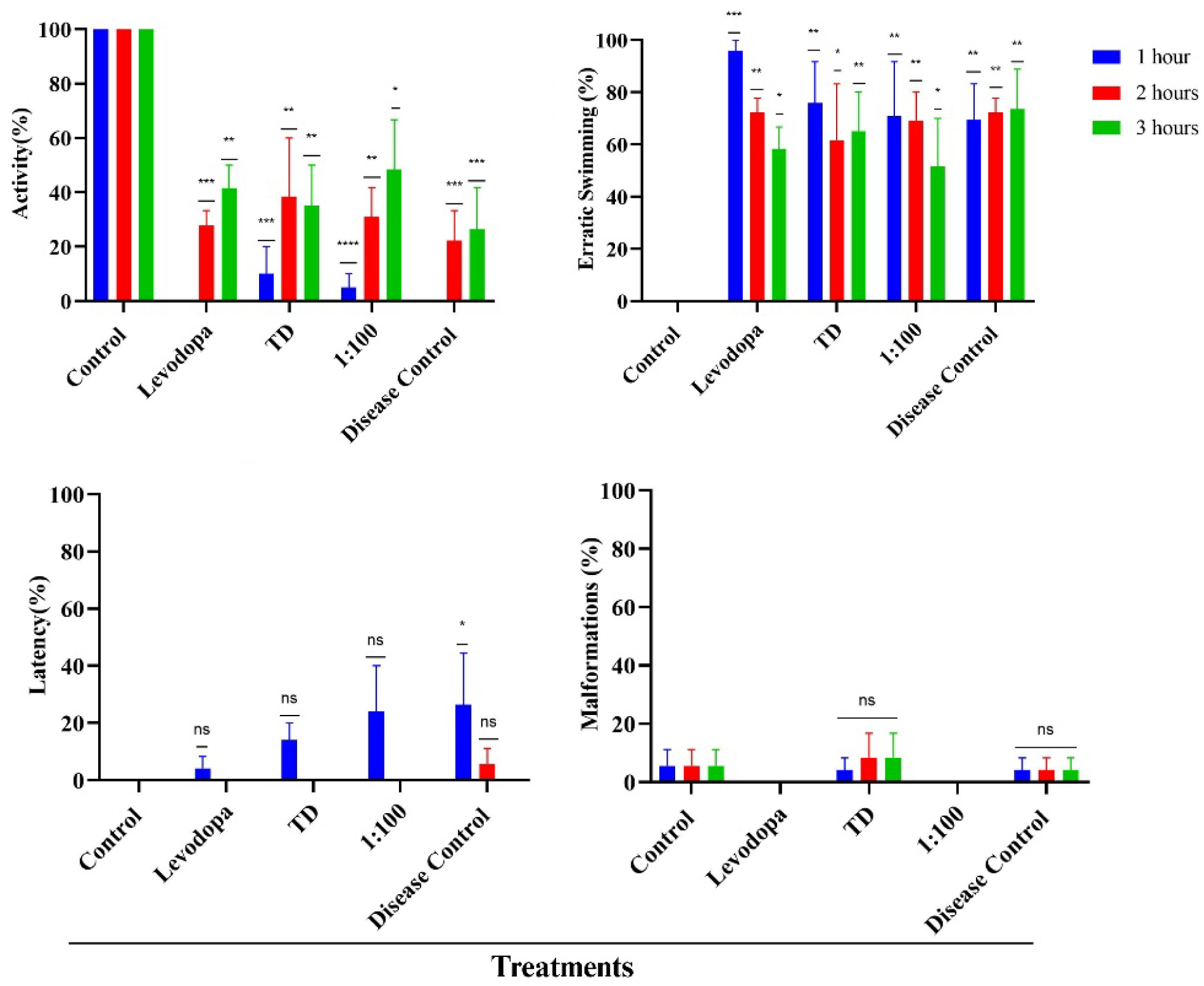
The morphological changes such as activity, erratic swimming, malformations and latency after inducing Parkinson’s disease and post-Parkinson’s disease treatments in 3dpf larvae with TD, Levodopa, TD:LD (1:100). The graph represents the various time points post-Parkinson’s disease treatments; 1 hr, 2hrs and 3 hrs with TD, Levodopa, TD:LD (1:100). (A) activity (B) erratic swimming (C) malformations (D) latency. Data represent means ± SEM of three independent experiments. ***Statistically significant p-value (p <0.001), **Statistically significant p-value (p <0.01), *Statistically significant p-value (p < 0.05) when compared with control.

The most evident change was observed in the TD:Levodopa (1:100) group, where locomotor activity increased from 30.8% at the first hour to 48.3% by the second hour, which indicates recovery in motor function. The free Levodopa and TD only group reached 41.6% and 35%, respectively, by the third hour. The TD:Levodopa group showed a consistently higher recovery than that observed in the free Levodopa and TD-only groups. In contrast, the disease control group showed lower activity, reaching only 26% at the end of the observation period. The observations suggest that the TD:Levodopa (1:100) group enhances therapeutic availability over time, which indicates a controlled or sustained drug release mechanism. This helps in maintaining the therapeutic effect of the drug for a longer time duration compared to the rapid release, which is associated with free drug treatment. These locomotor improvements were further reflected in the latency trends observed across the treatment groups. Latency was initially detected in all groups, with the TD:Levodopa (1:100) group exhibiting a value of 24.1%, free Levodopa and TD only exhibited 4.1% and 14.1% in the first hour, which diminished to 0 at 2 hours. Elevated latency was also observed in the disease control group (26.3%) and was there up to the second hour (5.5%), which shows no improvement in the absence of effective treatment. The gradual reduction in latency over time reflects an improvement in neuromotor functions, which suggests that there is recovery of dopaminergic signalling pathways.

The behavioural recovery patterns were strongly associated with the erratic swimming responses observed in the larvae. By the third hour of observation, erratic swimming behaviour in the TD:Levodopa (1:100) group declined to 51.66%, indicating a shift towards normalized swimming behaviour and improved neuromotor coordination. In contrast, the disease control group exhibited a progressive increase in erratic movement, rising from 69.44% at 1 hour to 73.61% at 3 hours, suggesting persistence of neurological dysfunction. The free Levodopa and TD-only groups displayed intermediate levels of erratic swimming at 58.33% and 65%, respectively, both of which remained higher than the TD:Levodopa (1:100) group. Notably, the free Levodopa group initially exhibited extremely high erratic swimming behaviour (95.8%), which subsequently decreased to 58.3% within two hours. This rapid change suggests an immediate and sudden effect, due to rapid drug diffusion and clearance due to the lack of a sustained release as opposed to the TD:Levodopa group.

These observations highlight that the TD:Levodopa conjugate formulation functions as a stabilizing carrier that regulates the release of Levodopa, and reduces the abrupt fluctuations in the system which can occur with free drug administration. The improved behavioural stability and sustained locomotor recovery observed in the TD:Levodopa (1:100) group suggest that it is an effective therapeutic outcome, since it facilitates the drug delivery at a lower concentration and in a sustained manner. There is no toxicity of the treatment revealed, since no mortality across any of the treatment groups was observed during the 3-hour observation period. This shows that the treatments were tolerated by the larvae and did not have any side effects. Since TDs are biocompatible and capable of neurorestoration, they serve as a potent delivery agent for Levodopa.

Tremor induction is the most characteristic symptom of Parkinsonism due to motor impairment. We manually quantified the number of tremors observed per minute in a larva at the end of the treatment at 3 hours. A significantly high level of tremors was observed in the Parkinson induced group (82 tremors/min), and that remained nearly unchanged in the disease control group at 78 tremors/min, indicating that there is very low to no recovery without a treatment regime. Notably, all treatment groups represented a measurable reduction in tremor frequency, indicating restoration of neural-motor functions due to the treatments. The TD 1:100 treatment group showed the most prominent reduction at 50 tremors/min, indicating a greater restoration of function and a move towards normalcy. The free levodopa and only TD group showed a reduction in tremor frequency as well (55 tremors/min and 58 tremors/min). However, the TD 1:100 group shows a better therapeutic effect compared to other treatment groups. At the end of the 3hour period, a noticeable difference is seen in the TD 1:100 group because it helps in sustained release in contrast to the rapid diffusion by the free drug, which alleviates the effect, but the effect doesn’t sustain for a longer period of time.

The heart rate was measured for larvae in treatment, as it is indicative of the physiological stress on a vital organ, the heart, post treatment. The heart rate was recorded at the end of the treatment period, at the 3-hour mark. Heart rate in the Parkinson’s induced group decreased significantly to 58.1%, which indicates that the heart rate is much less than normal, suggesting systemic stress following disease induction. Comparatively, the heart rate in the treatment group TD only was comparable to the control group (104.9%), indicating that there is no additional stress of the nanocages on the system. However, the heart rate was lower in the TD :Levodopa group at 94.1% and in the Levodopa group at 91.2%, indicating reduced stress on the larval system and restoration of normal physiological functions after treatment. These values are higher than those observed in the disease control group (80%), due to a lack of treatment. The TD:Levodopa group has shown a better and regulated response compared to the free Levodopa group. This suggests that there is an increase in treatment efficiency in delivering Levodopa with TD, and the prevalence of maintained homeostasis in the system.

**Figure 4:**
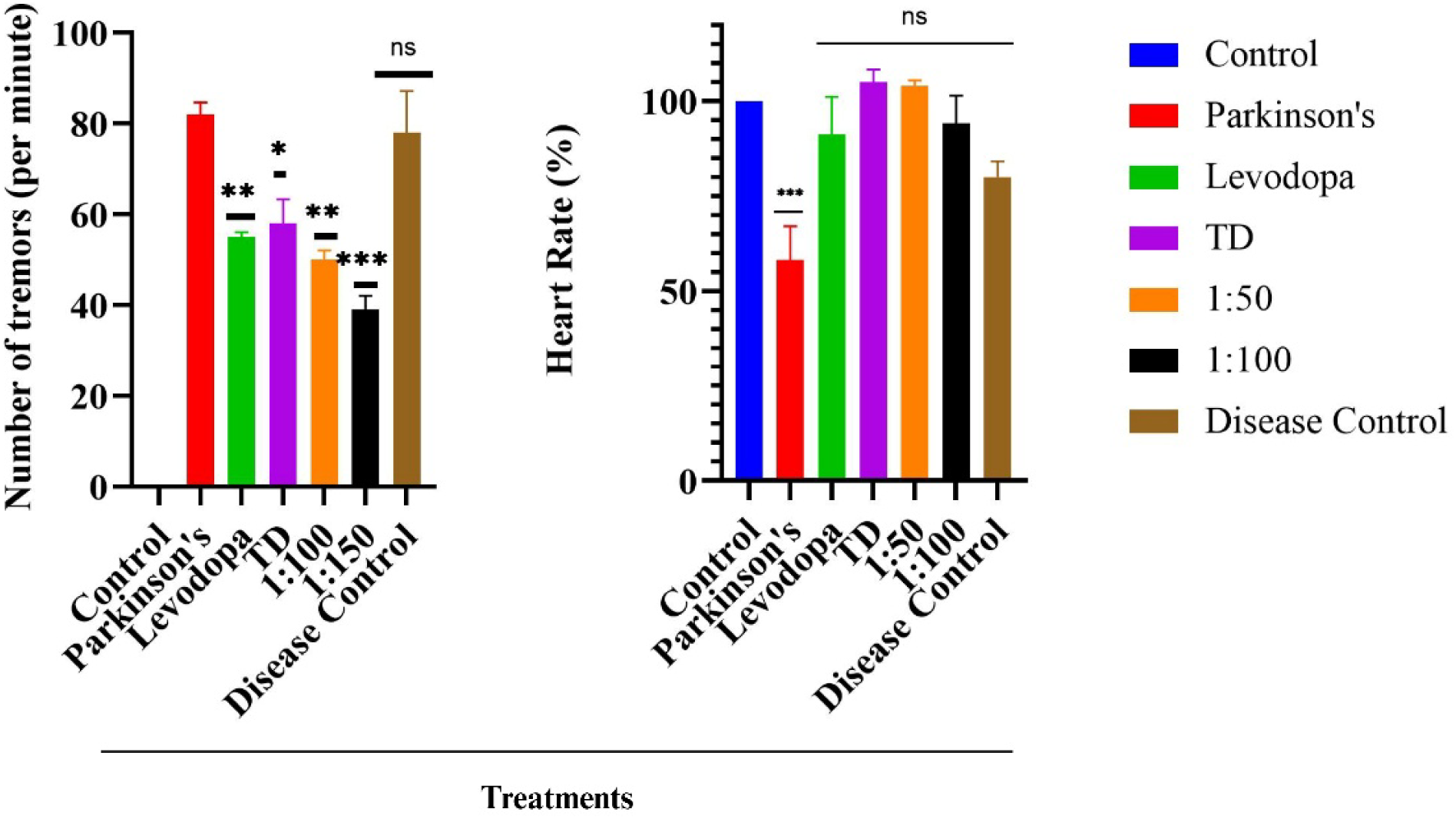
Tremors and heartbeat analysis after inducing Parkinson’s disease and post-Parkinson’s disease treatments in 3dpf larvae with TD, Levodopa, TD:LD (1:50) and (1:100). Data represent means ± SEM of three independent experiments. ***Statistically significant p-value (p <0.001), **Statistically significant p-value (p <0.01), *Statistically significant p-value (p < 0.05) when compared with control.

### Intracellular reactive oxygen species

DCFDA staining was used to evaluate the levels of intracellular reactive oxygen species, and the corresponding fluorescence intensity was quantified using ImageJ software. Accumulation of ROS is a crucial indicator of oxidative stress in a biological system. In this study, a marked elevation in ROS levels was observed in the Parkinson’s group, reaching a 2.1-fold increase relative to the control group. This substantial rise reflects the oxidative imbalance induced by Parkinsonian pathology and suggests considerable cellular stress and metabolic dysfunction following disease induction.

For the recovery groups, TD:Levodopa (1:100) and TD-only groups, marginally higher values 2.271-fold and 2.278-fold change, respectively were observed than those in the Parkinson’s group. The slightly elevated oxidative stress could be an indication of an increased metabolic activity associated with the recovery process. The similarity between the two treatments indicates the potential contribution of TD nanocages in stabilizing the oxidative environment within the system. On the contrary, free Levdopa treatment group showcased the highest ROS levels, reaching a 2.74-fold increase relative to the control. This elevated level indicates that even with the presence of a known therapeutic agent, the system experiences notable oxidative stress, possibly because of the rapid drug metabolism of the free drug. Similarly, the disease control group recorded ROS levels of 2.2-fold change, which remain comparable to the Parkinson’s group and further highlight the persistence of oxidative stress in the absence of effective therapeutic intervention. The presence of TD potentially facilitates improved cellular response and may contribute to the activation of antioxidant defence mechanisms and neurorestorative pathways, thereby supporting recovery in Parkinsonian zebrafish larvae.

**Figure 5:**
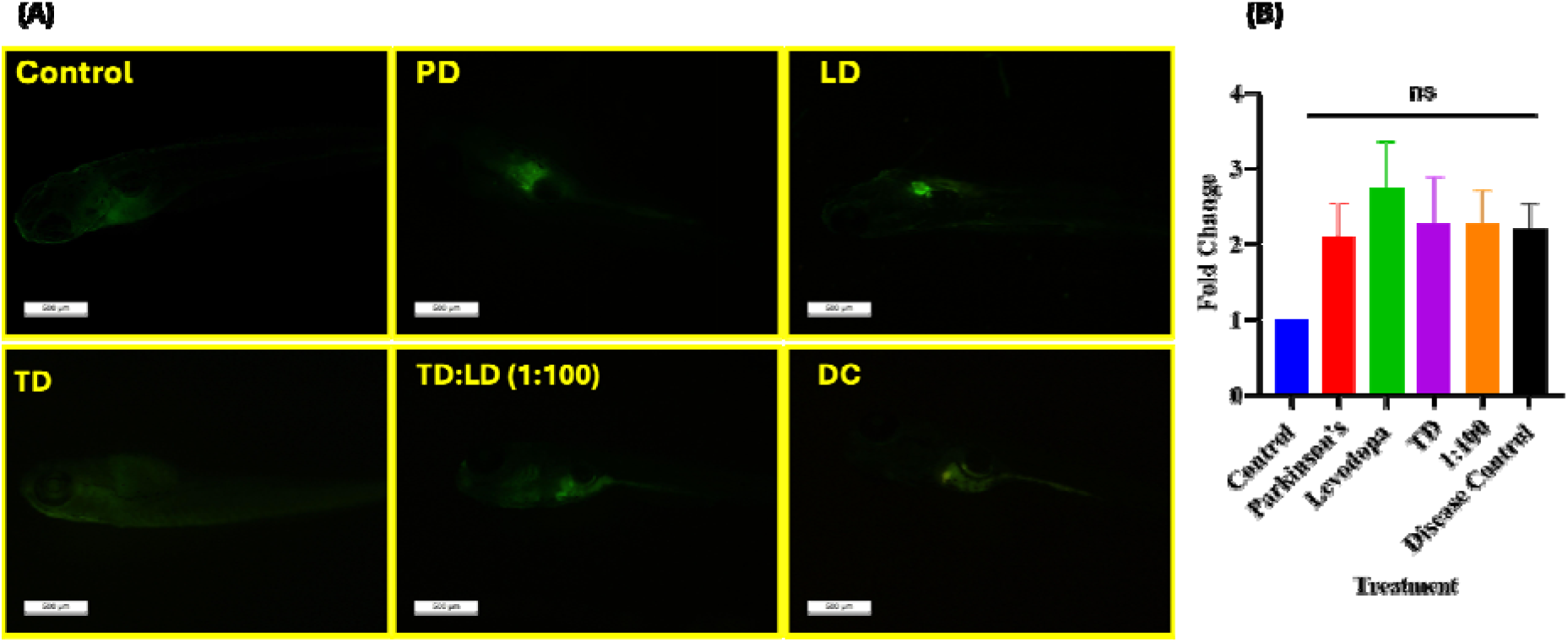
(A) Intracellular ROS generation detected by DCFDA dye. 3dpf larvae were exposed to TD, Levodopa, TD:LD (1:100) post Parkinson’s disease (B) Bar graph depicts fold change in ROS generation. Data represent means ± SEM of three independent experiments.

### Programmed Cell Death

Apoptosis is also known as programmed cell death and is closely associated with intracellular reactive oxygen species (ROS) levels. In the present study, the Parkinson’s group showed a high level of apoptosis, with levels reaching a 1.69-fold change relative to the control group. This indicates that there are impaired cellular functions associated with Parkinson’s disease, and also that neuronal cell death is occurring. Similarly, the disease control group showed an elevation of 1.64-fold change, which indicates that there is no improvement in the absence of a therapeutic regime, and cellular death continues to occur. In contrast, the larva treated with TD:Levodopa (1:100) showed a significant reduction in apoptosis at 0.72-fold change relative to the control group. The free Levodopa and TD-only group showed a 1.25- and 1.03-fold change increase, respectively, which are both higher than those observed in the control group. This suggests that Levodopa, when delivered with TD, helps in mitigating apoptosis in the system and restores normal functioning of the larvae post Parkinsonism.

**Figure 6:**
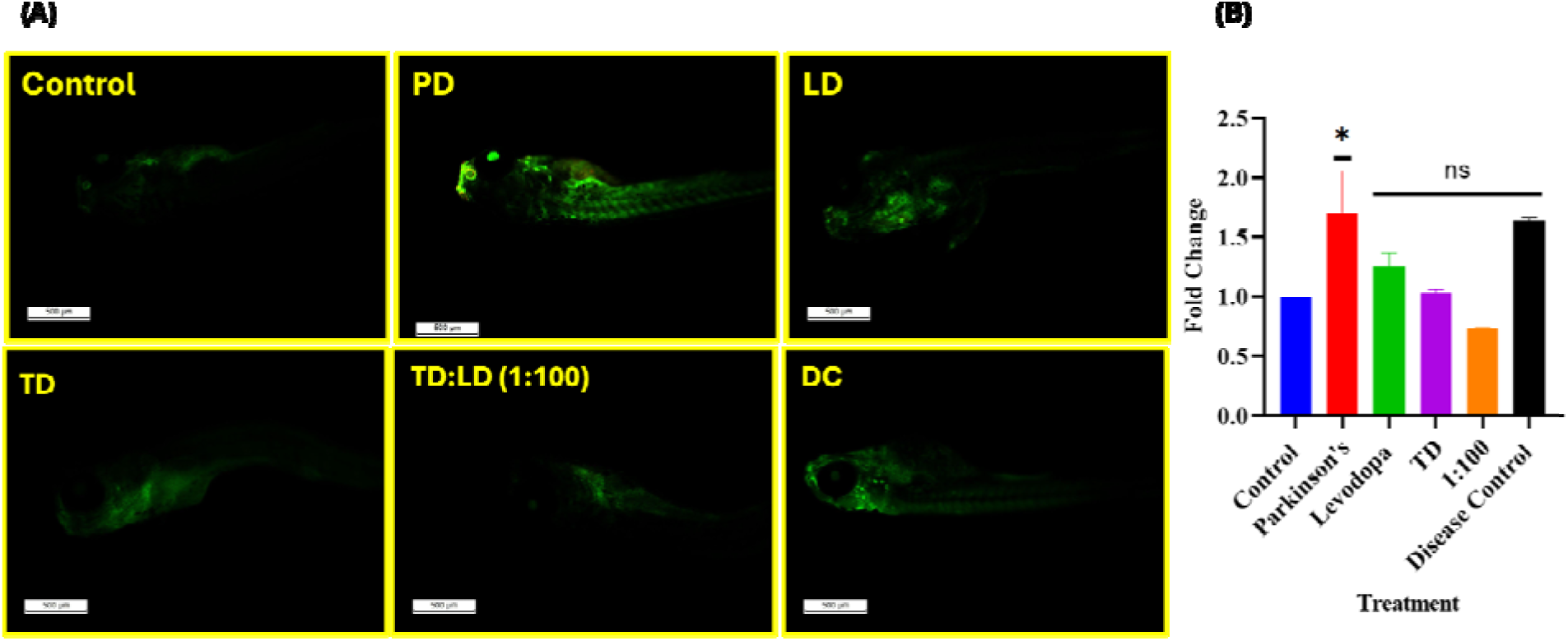
A programmed cell death detected by acridine orange. 3dpf larvae were exposed to TD, Levodopa, TD:LD (1:100) post Parkinson’s disease (B) Bar graph depicts fold change in ROS generation. Data represent means ± SEM of three independent experiments. *Statistically significant p-value (p < 0.05) when compared with control.

### Enzyme linked immunosorbent assay (ELISA) to quantify Dopamine

To assess the neuroprotective effectiveness of DNA nanocage-mediated levodopa administration against MPTP-induced Parkinsonian neurotoxicity, the dopamine (DA) content in zebrafish larvae was measured using ELISA (Figure 7). At roughly 13.5 pg/mL, the control group’s DA concentration was the highest, indicating normal dopaminergic neurotransmission. On the other hand, MPTP-treated larvae demonstrated a significant decrease in DA levels around 7.0 pg/mL, indicating that Parkinsonian disease was successfully induced through the selective degeneration of dopaminergic neurons. DA levels in the disease control group only partially recovered which was 10.0 pg/mL, staying below those of the healthy control, suggesting ongoing dopaminergic dysfunction. A moderate DA concentration around 8.5 pg/mL was obtained after treatment with TDs alone, indicating that although TDs are biocompatible, they do not significantly raise dopamine levels in the absence of a therapeutic agent. When free levodopa was administered, the concentration of dopamine rose to about 9.7 pg/mL, indicating its proven capacity to restore dopamine after MPTP-induced depletion. Remarkably, the TD:Levodopa (1:100) formulation also considerably raised DA levels, around 9.0 pg/mL, suggesting that levodopa was successfully delivered via the TDs. The nanocarrier-based formulation offers additional benefits, such as improved drug stability, enhanced cellular uptake, protection from premature degradation, and the potential for sustained drug release, which may contribute to prolonged therapeutic efficacy, even though the absolute dopamine concentration was comparable to that obtained with free levodopa.

**Figure 7:**
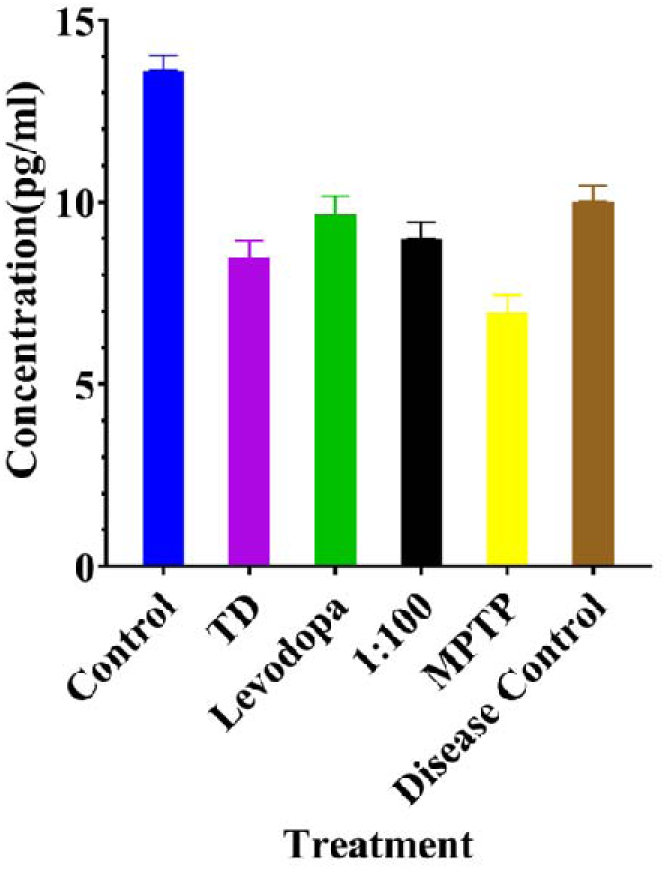
Quantification of dopamine (DA) levels in zebrafish larvae by ELISA. Dopamine concentrations were measured in the Control, TD, Levodopa, TD:Levodopa (1:100), MPTP, and Disease Control groups to evaluate the neuroprotective efficacy of DNA nanocage-mediated levodopa delivery in the MPTP-induced Parkinson’s disease model. Data represent means ± SEM of three independent experiments.

While the equivalent recovery attained by the TD:Levodopa formulation demonstrates the capacity of DNA nanocages to effectively deliver levodopa to target tissues, the restoration of dopamine levels after levodopa treatment verifies replenishment of the depleted neurotransmitter pool. DNA nanocages’ neuroprotective qualities, along with their superior biocompatibility and effective intracellular transport, may further lessen oxidative stress and maintain the integrity of dopaminergic neurons. The ELISA results are in line with the treated groups’ reduced oxidative stress, decreased apoptosis, elevated Park7/DJ-1 levels and behavioral recovery. All of these findings show that levodopa administration via DNA nanocage successfully reduces MPTP-induced dopamine depletion and maintains dopaminergic function in zebrafish larvae. The work emphasizes the potential of programmable DNA nanocages as a promising nanotherapeutic platform for improving Parkinson’s disease treatment results and levodopa administration.

### Detection of Park7/DJ-1 expression in Zebrafish Larvae by Immunofluorescent Assay

Significant differences in Park7/DJ-1 expression were shown by immunofluorescence analysis between the experimental groups, indicating modifications in the antioxidant defence system during Parkinsonian neurodegeneration. Park7/DJ-1 fluorescence intensity was lowest in the MPTP-treated group, suggesting that this neuroprotective protein was suppressed after MPTP-induced dopaminergic neuronal damage (Figure 8). DJ-1’s decreased expression indicates compromised cellular defense systems and heightened vulnerability to oxidative damage because it is essential for shielding neurons from oxidative stress and preserving mitochondrial integrity. In contrast, the basal levels of Park7/DJ-1 fluorescence in the control and tetrahedron (TD) groups were similar, indicating that the DNA tetrahedrons by themselves did not negatively impact endogenous DJ-1 expression and are biocompatible in physiological settings. When compared to the MPTP group, treatment with free levodopa dramatically increased Park7/DJ-1 expression, suggesting a partial restoration of the cellular antioxidant response. Among all study groups, zebrafish larvae exposed with the TD:LD (1:100) formulation showed the strongest Park7/DJ-1 fluorescence intensity, indicating better neuroprotection than free levodopa (Figure 8). DJ-1 expression in the disease control group only slightly increased, and it was still much lower than in the TD:LD-treated group. By scavenging reactive oxygen species, controlling mitochondrial homeostasis, and averting oxidative stress-induced neuronal death, the multifunctional redox-sensitive protein DJ-1 (Park7) safeguards dopaminergic neurons.

**Figure 8:**
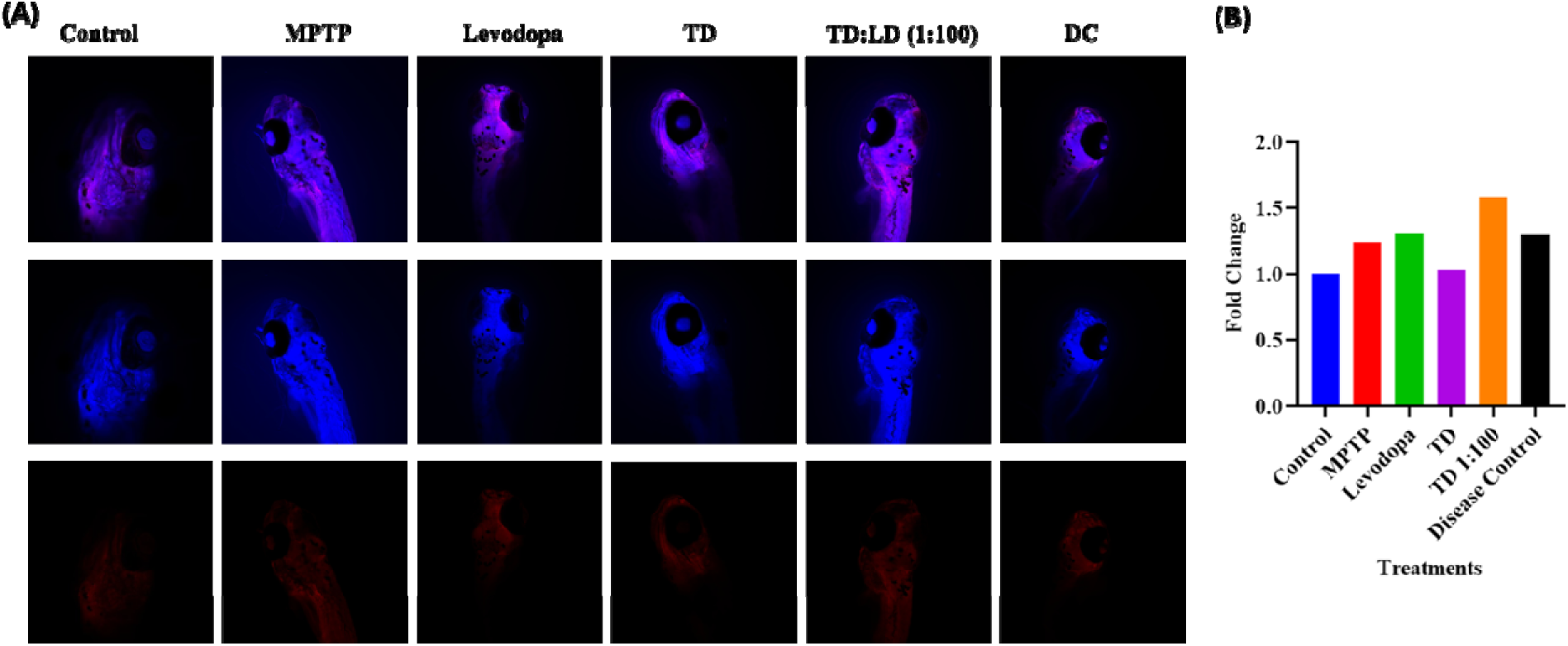
Quantitative analysis of Park7/DJ-1 expression in Parkinson’s induced 3 dpf zebrafish larvae by immunofluorescence assay. Fluorescence intensity of Park7/DJ-1 was quantified in the Control, DNA tetrahedron (TD), Levodopa, TD:Levodopa (1:100), MPTP, and Disease Control groups. (A) Representative photographs. (B) Bar graph depicts quantitative analysis.

DJ-1 plays a crucial part in the pathophysiology of Parkinson’s disease, as evidenced by the substantial correlation between mutations or decreased expression of the protein and both familial and sporadic forms of the disease. Degeneration of dopaminergic neurons, excessive formation of reactive oxygen species, and mitochondrial dysfunction are caused by MPTP’s metabolism to MPP⁺, which preferentially inhibits mitochondrial complex I. Therefore, high oxidative stress and weakened endogenous antioxidant defences are consistent with the significant decrease in Park7/DJ-1 expression seen in MPTP-treated larvae. Park7/DJ-1 is significantly upregulated after TD:LD (1:100) treatment, indicating that DNA tetrahedron-mediated levodopa delivery restores neuronal antioxidant capacity more successfully than free levodopa alone (Figure 8). DNA tetrahedrons’ superior biocompatibility, stronger cellular absorption, and prolonged intracellular delivery, which lead to better maintenance of mitochondrial function and activation of endogenous neuroprotective pathways, are probably responsible for the increased therapeutic efficiency. These results corroborate the TD:LD (1:100) nanoplatform’s capacity to reduce oxidative stress and shield dopaminergic neurons from MPTP-induced neurotoxicity. They are also in line with improvements seen in other behavioral and biochemical markers. When taken as a whole, the immunofluorescence data show that DNA tetrahedron-assisted levodopa administration greatly increases Park7/DJ-1 expression, indicating its potential as a promising nanotherapeutic approach for enhancing neuroprotection and delaying the advancement of Parkinson’s disease.

### DNA Nanocage-Mediated Levodopa Delivery Restores Parkinson’s Disease-Associated Gene Expression

#### Genes associated with PD

The tyrosine hydroxylase (TH) gene is responsible for encoding a rate-limiting enzyme for the conversion of tyrosine to L-DOPA, a dopamine precursor. It is mainly expressed in dopaminergic neurons and regulates motor control. Degeneration of these neurons in PD results in reduced expression of TH and dopamine depletion. TH levels are commonly used as biomarkers to evaluate neuronal loss and disease progression in PD model. Depletion in TH levels is linked with reduced dopamine, along with motor dysfunction[31, 32]. In the present study, TH expression was observed to be slightly reduced in the Parkinson’s group, with a fold change of 0.87 relative to the control. A reduction was observed in the free Levodopa treatment group, where TH expression dropped substantially to 0.35-fold change, suggesting that administration of the free drug does not effectively restore endogenous dopamine synthesis pathways. In contrast, the TD:Levodopa (1:100) group exhibited a comparatively higher expression level of 0.47-fold change, indicating partial restoration of TH expression. The TD-only group showed a fold change of 0.83, which is comparable to the Parkinson’s group and suggests that the nanostructure itself does not adversely affect dopaminergic gene expression. Interestingly, the disease control group displayed a TH expression level of 0.41-fold change, which remains substantially lower than the control, indicating presence of dopaminergic dysfunction in the absence of an effective treatment regime (Figure 9).

**Figure 9:**
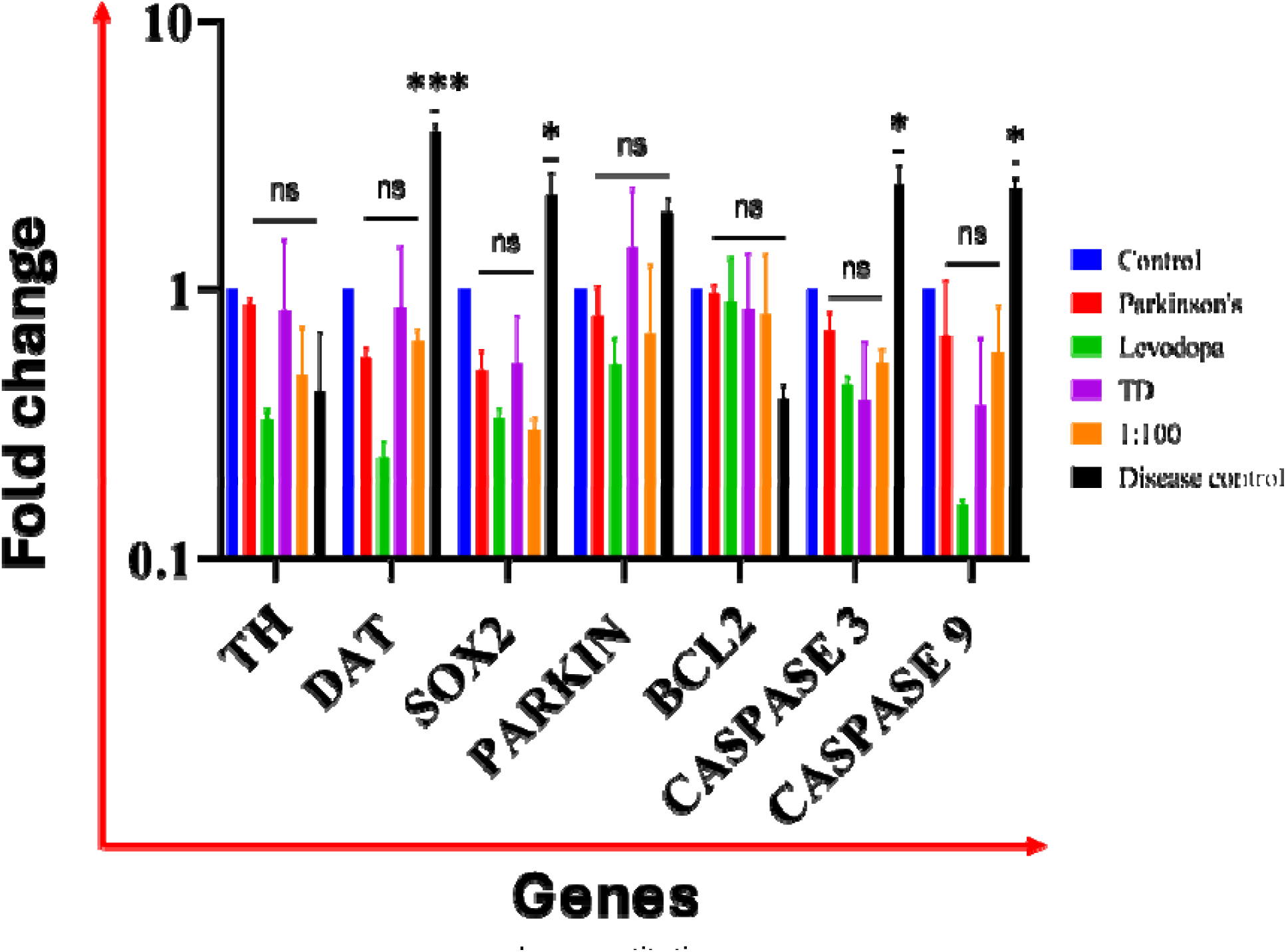
Relative expression of Parkinson’s disease-associated genes in zebrafish larvae determined by quantitative real-time PCR. Gene expression was analyzed in Control, TD, Levodopa, TD:Levodopa (1:100), MPTP, and Disease Control groups. Relative transcript levels were normalized to the housekeeping gene (β-actin/GAPDH) using the 2^−ΔΔCt method. Data are presented as mean ± SEM (n = 3). Statistical significance was determined by one-way ANOVA. ***Statistically significant p-value (p <0.001), **Statistically significant p-value (p <0.01), *Statistically significant p-value (p < 0.05) when compared with control.

DAT (Dopamine Transporter, SLC6A3) is responsible for the production of a membrane protein that regulates dopamine reuptake from the synaptic cleft into presynaptic neurons. Reduced DAT expression is observed in PD models, leading to impaired dopamine homeostasis[33]. Consistent with this phenomenon, DAT expression in the Parkinson’s group was markedly reduced to 0.55-fold change relative to the control group. In contrast, the disease control group demonstrated an abnormally elevated expression of 3.8-fold change, indicating highly dysregulated gene activity and abnormal dopamine transporter regulation under diseased conditions without treatment. The free Levodopa treatment group showed a significantly reduced DAT expression of 0.23-fold change, suggesting that dopamine alone cannot restore the gene regulation. Conversely, the TD-only and TD:Levodopa (1:100) treatment group indicated expression levels closer to control, with fold change of 0.85 and 0.63, respectively. This indicates that TD-mediated delivery of Levodopa could potentially stabilize dopamine delivery mechanism and a partial restoration of dopamine homeostasis (Figure 9).

Sox2 encodes a transcription factor crucial for maintaining pluripotency and self-renewal of neural stem cells. It controls neural lineage commitment and stem cell differentiation to neurons, even dopaminergic neuron precursors. In PD models, SOX2 modulation facilitates the generation of dopaminergic neurons and supports neural regeneration, thereby serving as a key factor in neural repair[34]. In this study, SOX2 expression in the Parkinson’s group was significantly reduced to 0.49-fold change relative to the control group, indicating impaired neural regeneration under Parkinsonian conditions. The free Levodopa treatment group showed a reduction in SOX2 expression to 0.33-fold change, suggesting limited activation of regenerative pathways following treatment with the free drug. Similarly, the TD:Levodopa (1:100) group demonstrated a fold change of 0.29, indicating that SOX2-mediated regeneration may not be fully restored within the observed treatment window. The TD-only group showed a fold change of 0.52, which is comparable to the Parkinson’s group and suggests that the nanostructure does not significantly alter the gene’s activity. Notably, the disease control group displayed an increase in SOX2 expression to 2.2-fold change, indicating a strong but potentially dysregulated activation of regenerative pathways under persistent disease stress.

Parkin encoded by PARK2 gene is a E3 ubiquitin ligase. It is responsible for tagging damaged or misfolded proteins for further degradation via ubiquitin-proteasome system, thereby maintaining mitochondrial quality control through mitophagy. In cases of PD, loss or dysfunction of parkin can lead to accumulation of damaged mitochondria and protein aggregates. Reduced parkin functions disrupt mitophagy, protein clearance, increasing oxidative stress and mitochondrial dysfunction[35]. In the present study, Parkin expression in the Parkinson’s group was reduced to 0.79-fold change relative to the control group, indicating impaired mitochondrial maintenance mechanisms under disease conditions. In the free Levodopa treatment group, Parkin expression further declined to 0.51-fold change, suggesting limited recovery of mitochondrial maintenance processes (Figure 9). In contrast, the TD-only group showed an increased expression level of 1.4-fold change, indicating enhanced activation of protective mitochondrial maintenance pathways. The TD:Levodopa (1:100) group displayed a fold change of 0.67, suggesting partial restoration of Parkin activity. Interestingly, the disease control group demonstrated an abnormally elevated expression level of 1.9-fold change, indicating excessive activation of stress response pathways and a deviation from normal cellular regulation in the absence of therapeutic intervention.

### Genes associated with apoptosis

Bcl2 and Bcl-xL, anti-apoptotic proteins maintains the integrity of outer mitochondrial membrane (OMM) and inhibits the release of cytochrome C, thereby blocking the apoptotic pathway activation[36]. Elevated calcium levels can trigger intrinsic apoptotic pathway through calpain activation and inducing the opening of the mitochondrial permeability transition pore. Activated calpains and caspase 8 cleaves Bid, thereby producing truncated Bid, which can translocate to mitochondria, and counteract the anti-apoptotic proteins and interacts with pro-apoptotic proteins. Consequently, the mitochondrial permeability transition pore opens, leading to the release of cytochrome C and other factors into the cytoplasm. In the presence of dATP, an apoptosome is formed comprising cytochrome C, Apaf-1, and procaspase-9. This apoptosome further activates caspase-9, which subsequently triggers the downstream activation of caspase-3. Generally, elevated levels of anti-apoptotic genes are associated with reduced expression of pro-apoptotic genes. Numerous studies have demonstrated that ROS-induced cell death in zebrafish can be analysed through the expression levels of apoptotic genes[27]. In this study, the expression levels of BCL2 in the treatment groups were relatively comparable. The free Levodopa group exhibited a fold change of 0.90, while the TD-only group and TD:Levodopa (1:100) group showed fold changes of 0.84 and 0.80 respectively. These values are close to the control levels, suggesting that the apoptotic pathway is being effectively moderated in these treatment conditions. In contrast, the disease control group displayed a marked reduction in BCL2 expression to 0.39-fold change, indicating a loss of anti-apoptotic protection and increased neuronal cell death. This observation is further supported by the expression patterns of the pro-apoptotic genes CASPASE-3 and CASPASE-9. In the present study, CASPASE-3 expression in the Parkinson’s group was measured at 0.70-fold change relative to the control. Notably, all treatment groups demonstrated a reduction in CASPASE-3 expression compared to the Parkinson’s group. A value of 0.43-fold change was observed in the free Levodopa treatment group, while the TD-only group showed a value of 0.38. The TD:Levodopa (1:100) group showed a fold change of 0.52, which indicates partial suppression of apoptotic signalling (Figure 9). These observations suggest that the treatments are associated with mitigation of apoptosis in the system. In contrast, the disease control group exhibited a dramatic increase in CASPASE-3 expression to 2.4-fold change, indicating active apoptotic processes and neuronal degeneration in the absence of therapeutic intervention. In this study, CASPASE-9 expression in the Parkinson’s group was measured at 0.66-fold change relative to the control group. The free Levodopa treatment group displayed an extremely low expression level of 0.16-fold change, which may indicate excessive suppression of apoptotic signalling and potential disruption of normal cellular metabolic processes. The TD-only group exhibited a reduced expression level of 0.36-fold change. Conversely, the TD:Levodopa (1:100) group showed a moderate 0.58-fold change in expression, suggesting that apoptotic activity is moderated without completely disrupting normal cellular and metabolic processes. In contrast, the disease control group showed an elevated expression of 2.36-fold change, indicating extensive activation of apoptotic pathways and severe cellular stress in the absence of therapeutic treatment.

## Conclusions

In this study, we used an MPTP-induced zebrafish larval model to design and assess a DNA tetrahedron (TD)-mediated levodopa delivery system as a nanotherapeutic approach for the treatment of Parkinson’s disease. Levodopa was effectively delivered using the DNA nanocage as a biocompatible and programmable carrier, maintaining the medication’s therapeutic effectiveness. The TD:Levodopa (1:100) combination had the most promising therapeutic efficacy among the evaluated formulations, indicating its potential as an optimized nanomedicine for Parkinson’s disease. Key pathological features of Parkinson’s disease, such as decreased locomotor activity, tremors, unpredictable swimming behavior, increased latency, elevated oxidative stress, apoptosis, dopamine depletion, and dysregulation of genes related to dopaminergic neurotransmission, neuronal survival, and mitochondrial homeostasis, were successfully replicated by MPTP exposure. These pathogenic changes validate the zebrafish model as an appropriate platform for treatment investigation since they closely match the gradual degradation of dopaminergic neurons seen in human Parkinson’s disease.

These Parkinsonian symptoms were successfully reduced at several biological levels by TD-mediated levodopa treatment. Improvements in swimming behavior and locomotor performance demonstrated behavioral recovery and the restoration of neural function. Additionally, biochemical investigations showed that the TD:Levodopa formulation effectively attenuated oxidative stress-mediated neuronal injury by dramatically restoring dopamine levels while lowering intracellular reactive oxygen species and apoptotic cell death. Significantly, real time PCR analysis showed normalization of genes linked to dopamine synthesis and transport (TH and DAT), neuronal differentiation and regeneration (SOX2), mitochondrial quality control (PARKIN), and apoptosis (BCL2 and caspases), while immunofluorescence analysis showed restoration of neuroprotective markers. Together, our results suggest that levodopa administration via DNA nanocages produces neuroprotective benefits by coordinating the control of several physiological pathways instead than only replenishing dopamine. The DNA nanocage platform’s special physicochemical characteristics; such as superior biocompatibility, structural programmability, effective cellular internalization, protection of the encapsulated drug from premature degradation, and the possibility of controlled and prolonged drug release; are responsible for its therapeutic efficacy. DNA tetrahedron-mediated delivery provides an advanced nanocarrier platform that can improve therapeutic performance while reducing systemic drug loss, in contrast to traditional levodopa therapy, which is frequently linked to limited bioavailability, fluctuating plasma concentrations, and long-term motor complications. These features make DNA nanocages very appealing for enhancing the accuracy and effectiveness of medication delivery to the neurological system.

Overall, this study offers thorough proof that levodopa delivery via DNA tetrahedron dramatically reduces Parkinsonian symptoms by reestablishing molecular pathways necessary for neuronal survival, reducing oxidative stress and apoptosis, and restoring dopaminergic function. To the best of our knowledge, this is one of the first studies to show how DNA tetrahedron nanocages may be used to distribute levodopa specifically in an in vivo Parkinson’s disease model. These results establish DNA nanocages as a promising platform for nanomedicine-based therapies in neurodegenerative illnesses and broaden the biological uses of structured DNA nanotechnology beyond traditional drug delivery. Prior to clinical translation, more research in mammalian models is necessary to assess pharmacokinetics, biodistribution, blood-brain barrier transit, long-term biocompatibility, and therapeutic efficacy, even if the current work demonstrates proof-of-concept in a zebrafish model. Treatment results may be further improved by future research using targeted surface modifications and multifunctional DNA nanocages that can co-deliver neuroprotective drugs, antioxidants, or gene therapies. Together, these efforts provide a promising substitute for traditional medication delivery methods by laying the groundwork for the creation of next-generation DNA nanotechnology-based precision treatments for Parkinson’s disease and other neurodegenerative diseases.

## Acknowledgements

The authors would like to express their sincere gratitude to all members of the A.K. and D.B. groups for their critical evaluation of the work and insightful comments. K.K. expresses gratitude to GoI and DST for the National Postdoctoral Fellowship in Nanoscience and Technology. D.B. acknowledges research funds from ANRF-ARG, GSBTM; MoES for the STARS award; IITGN for the startup grant; and SERB and GoI for the Core research grant. A.K. thanks the GSBTM (GSBTM/JD(R&D)/663/2023-24/02003699) and ICMR (IIRP-2023-1078) for their financial assistance.

## Author Contributions

K.K. conceived the idea, planned all the experiments, analysed the results and wrote the original manuscript. S.S. and B.P. executed AFM, DLS and Confocal image analysis. S.C. and K.J. carried out *in vivo* experiments in zebrafish. A.K. provided the zebrafish facility and required funding. D.B. contributed the facility and required funding. All the authors discussed the data and helped in writing and critically reading the manuscript.

